# Accurate Detection of Tandem Repeats from Error-Prone Sequences with EquiRep

**DOI:** 10.1101/2024.11.05.621953

**Authors:** Zhezheng Song, Tasfia Zahin, Xiang Li, Mingfu Shao

**Author notes:** contribute equally to this work (Zhezheng Song, Tasfia Zahin).

## Abstract

A tandem repeat is a sequence of nucleotides that occurs as multiple contiguous and near-identical copies positioned next to each other. These repeats play critical roles in genetic diversity, gene regulation, and are strongly linked to various neurological and developmental disorders. While several methods exist for detecting tandem repeats, they often exhibit low accuracy when the repeat unit length increases or the number of copies is low. Furthermore, methods capable of handling highly mutated sequences remain scarce, highlighting a significant opportunity for improvement. We introduce EquiRep, a tool for accurate detection of tandem repeats from erroneous sequences. EquiRep estimates the likelihood of positions originating from the same position in the unit by self-alignment followed by a novel approach that refines the estimation. The built equivalent classes and the consecutive position information will be then used to build a weighted graph, and the cycle in this graph with maximum bottleneck weight while covering most nucleotide positions will be identified to reconstruct the repeat unit. We test EquiRep on simulated and real HOR and RCA datasets where it consistently outperforms or is comparable to state-of-the-art methods. EquiRep is robust to sequencing errors, and is able to make better predictions for long units and low frequencies which underscores its broad usability for studying tandem repeats.

## 1 Introduction

The human genome consists of a vast array of repetitive elements, and many of them arise from a process called tandem duplication. In this process, a segment of the DNA is replicated multiple times, creating consecutive approximate repeat units. The length of these repeat units vary from a few base pairs (called short tandem repeats or STRs) to a hundred base pairs (called variable number tandem repeats or VNTRs) and sometimes upto thousand base pairs in satellite DNAs. Tandem repeats make up about 8-10% of the human genome and have been closely linked to several neurological and developmental disorders like Huntington’s disease, Friedreich’s Ataxia, fragile X syndrome, etc [7,18,20]. The repeat tracks associated with many of these diseases appear longer in certain affected individuals than typically observed in the general population [7,18,20]. For example, the GAA unit associated with Friedreich’s Ataxia appears 5-30 times normally, but 66 to over 1000 times in affected individuals [2]. More recently, longer repeats copies (25-30bp) have been discovered to influence schizophrenia [19] and Alzheimer’s disease [3]. Alpha satellite repeats of about 171 bp organized into Higher Order Repeat (HOR) units are found to be abundant in centromeric regions of many organisms and are essential for studying genome stability and evolutionary dynamics [12,13]. To analyze tandem repeats, a critical step often involves the accurate reconstruction of the unit from either assembled genome or unassembled (long) reads.

The Rolling Circle Amplification (RCA) is a recently refined sequencing technique that amplifies circularized template molecules, producing numerous tandem repeat copies of the original template. RCA can yield long tandem repeat units, with sequences often exceeding 150 bp and even reaching several kilobases in certain contexts. RCA followed by PacBio or Nanopore sequencing is a popular protocol adopted in many recent studies, specially for detection of full-length circular RNAs [23,24,11]. A crucial step in this process is the prediction of a consensus sequence derived from long reads, providing a highly accurate reconstruction of the repeat unit compared to the original reads with sequencing errors. This step requires *in silico* intervention, and typically employs widely used tandem repeat detection tools for consensus sequence prediction. It is important to emphasize that the reliability of circular RNA detection is therefore significantly influenced by the accuracy of the predicted consensus sequence during this intermediate step. Consequently, there is a pressing need for reliable tools capable of accurately predicting tandem repeat patterns of different kinds, accounting for the variability in unit length and copy number that may exist in different biological contexts. Addressing this gap is particularly essential for improving the accuracy and reliability of full-length circular RNA identification, especially considering that circular RNAs have emerged as promising biomarkers for numerous diseases [17,10,21].

Both above critical applications can be abstracted as this computational problem: given a sequence *R*, decide if *R* contains tandem repeats, and if yes, construct the unit. Many methods have been developed, mainly driven by the development of sequencing technologies. Tools include mreps [9], RepeatMasker (https://www.repeatmasker.org/), INVERTER [22] are primarily designed to detect small repeat units from relatively low error rate data such as short-read sequencing data. They often do not perform well with higher repeat lengths and/or lower frequencies. Other tools like DeepRepeat [5], tandem-genotypes [14], ExpansionHunter [4] emphasize more the quantification of tandem repeats than detection. Tandem Repeat Finder (TRF) [1] is one of the most widely used tandem repeat detection tools. It is based on the idea of k-tuple matching and utilizes a probabilistic model followed by statistical analysis to make repeat predictions. It is also suitable for use in erroneous long reads given its ability to handle substitutions and indels. With the advent of third-generation sequencing and the resulting access to long-reads data, new tools such as TideHunter [6] and mTR [15] began to emerge. TideHunter is an efficient tandem repeat detection and consensus calling tool tailored for RCA based long reads sequences. However, it faces challenges in accuracy when dealing with repeat of small length. Similarly, mTR struggles with repeats of low copy numbers, mostly due to difficulty in finding a long cycle of short, infrequent kmers. Despite the promising potential of long-reads in revealing novel disease-associated tandem repeats and in reconstructing full-length circRNAs, tools capable of managing high error rates are rare. Those currently available also struggle to achieve satisfactory accuracy in challenging settings (such as too short/long units and low copy numbers), as suggested by our experiments. Therefore, the task of accurately detecting tandem repeats from noisy sequences, particularly for longer units and low copy numbers, remains largely unresolved.

Here we present EquiRep, a new tool for detecting tandem repeats in error-prone sequences. EquiRep stands out for its robustness against sequencing errors, as well as its effectiveness in detecting repeats of low copy numbers. EquiRep employs a novel idea that identifies *equivalent* positions in the given sequence. This is achieved by self-local alignment followed by a critical refinement step that reduces the noises. The refined, equivalent positions are organized into equivalent classes. A graph is constructed where nodes are equivalent classes and the identification of unit can be formulated as searching for a cycle in the graph with maximized bottleneck weight. We evaluate the accuracy of EquiRep compared to 4 leading methods across 3 types of data. EquiRep drastically outperforms other tools on a diverse set of simulated data and a HOR dataset, while performing nearly the same with the best method on RCA data.

## 2 Methods

Given an error-prone (long) sequence/read *R*, EquiRep employs a 4-step approach to determine the true repeating unit *U* in it (if any).

**Step 1: identifying substring** *S* **with repeating structure**. Given the long read *R*, we first determine if *R* contains a substring *S* that potentially consists of multiple (mutated) tandem repeats. First, we use a classical seed-chaining procedure for a coarse-grained analysis. All kmers within the given long read are collected as seeds, and exact seed matches are identified to serve as anchors. EquiRep then applies a dynamic programming algorithm to group colinear anchors into chains, ultimately selecting the highest-scoring chain as the candidate substring *R*′.

*R*′ may not be accurate especially when the error rate is high which breaks shared kmers. We design a fine-grained approach aiming for more accurate detection. The technique we use here is the so-called *diagonal-free self-alignment*. That is, we perform local alignment between *R*′ and itself, but enforce that the same position cannot be aligned to itself. See Figure 1. This approach is efficient in determining the repeating structures. If *R*′ contains a repeating region, say with *m* units, then the optimal local alignment is likely made of the alignments between the *r*-th unit to the (*r* + 1)-th unit, 1 ≤ *r* ≤ *m* − 1. If *R*′ does not contain a repeating region, then the optimal local alignment often consists of short alignment with an insignificant alignment score. EquiRep performs the diagonal-free self-alignment on *R*′, and reports the repeating region *S*, which can be obtained by tracking back from the optimal solution if the optimal alignment length reaches 20.

**Fig 1:**
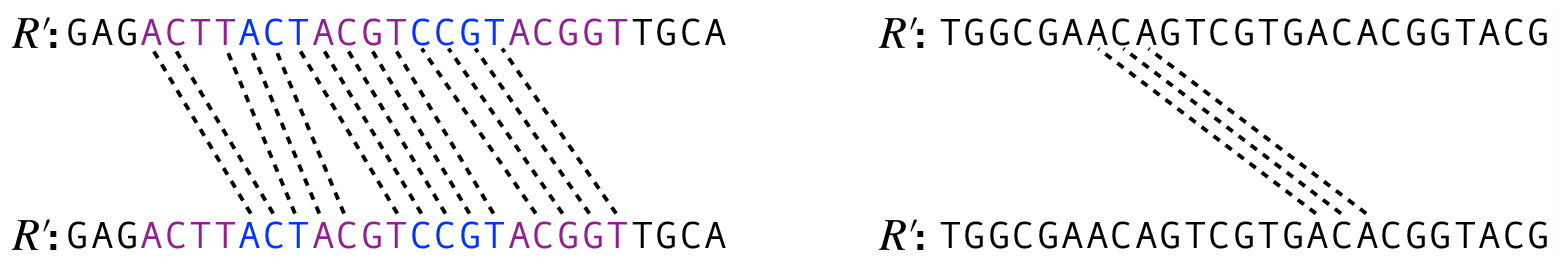
Left: *R′* contains a repeating region *S* = ACGTACTACGTCCGTACGGT. The true, unknown repeating unit *U* = ACGT, and *S* consists of *m* = 5 mutated such units. The optimal local alignment, given by dashed lines, aligns the *r*-th unit to the (*r* + 1)-th unit, *r* = 1, 2, 3, 4. Right: *R′* does not contain a repeating region.

**Step 2: constructing classes of equivalent positions 𝒞**. The core part of EquiRep is the construction of *equivalent positions*. See Figure 2. Formally, we define positions *i* and^3^*j* in *S* are equivalent, 1 ≤ *i < j* ≤ |*S*|, denoted as *i* ∼ *j*, if *S*[*i*] and *S*[*j*] originate from the same letter in the true unit *U*. For examples, in the left panel of Figure 1 we have 4 ∼ 8, 7 ∼ 10, etc. In ideal situations, the two positions in each aligned pair of the optimal diagonal-free self-alignment are equivalent, but in real situations, especially when the error rate is high, the optimal alignments are often erroneous, and therefore just using aligned pairs in the optimal alignment results in poor accuracy.

**Fig 2:**
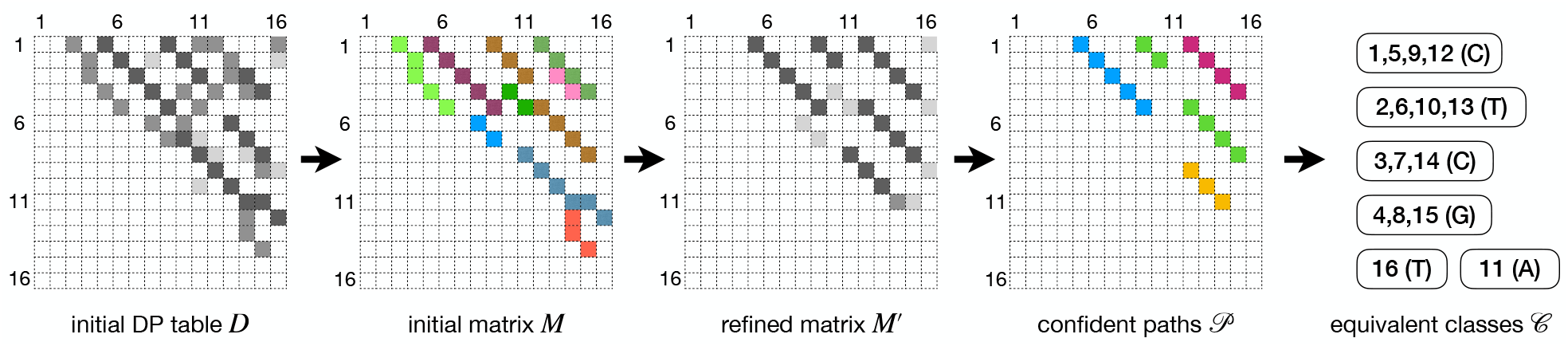
Illustration of constructing the equivalent classes 𝒞.

We propose a novel algorithm to accurately build equivalent positions. We start with the dynamic programming table *D* obtained directly from the diagonal-free self-alignment on *S*. This initial table *D* consists of rich, but noisy information about the repeating structures. The alignment between the *r*-th unit and the *s*-th unit, for all 1 ≤ *r < s* ≤ *m*, likely leads to significant alignment scores in table *D*. The alignments that correspond to the same *s* − *r* are also likely connected, forming significant paths in *D*. All these paths, not only the optimal path that corresponds to *s* − *r* = 1, are informative in constructing the equivalent positions. To collect these paths from *D*, we locate all entries in *D* that are local maxima within a 7 × 7 window (where 7 is a parameter of EquiRep) centered on each entry; we denote by *X* the set of entries that are local maxima. Entries in *X* are potential ending locations of significant paths. We process these entries in descending order of their alignment scores (i.e., *D*[, ]) to construct an initial matrix *M* of the size |*S*| × |*S*|. See Figure 2. Entries in *M* are all initialized as 0. For each entry (*p, q*) in *X*, we trace-back from it in *D* to retrieve a path *x*: the tracing-back terminates either when meeting an entry (*k, l*) such that *M* [*k, l*] ≠ 0, which means (*k, l*) is on one of the previously identified paths, or meeting an entry (*k, l*) with *D*[*k, l*] = 0, which marks the starting location of local alignment. In either case, the score of path *x* is set as *D*[*p, q*] − *D*[*k, l*]. If this score is below 25 (another parameter in EquiRep), we discard *x* as they typically result from random matchings; otherwise, we assign score *D*[*p, q*] − *D*[*k, l*] to *M* [*i, j*] for every entry (*i, j*) on the path *x*. At this time point, the equivalent information is encapsulated in *M* such that *M* [*i*][*j*] represents a loose estimation proportional to the likelihood that *i* ∼ *j*. But *M* is highly noisy, requiring further “refinement”.

We design a novel, efficient refinement algorithm based on this fundamental idea: for a tuple (*i, j, k*), if all three entries *M* [*i*][*j*], *M* [*i*][*k*], and *M* [*j*][*k*] are large, it indicates that the three pairs support each other and hence there is a higher probability that *i* ∼ *j, j* ∼ *k*, and *i* ∼ *k*. Based on this key observation, we design an iterative algorithm that refines *M*. In each round, we enhance all three entries for a tuple (*i, j, k*) by adding min(*M* [*i*][*j*], *M* [*i*][*k*], *M* [*j*][*k*]), followed by a row-sum normalization. EquiRep performs 5 rounds of refinements. The resulting refined matrix is stored to *M* ′. The pseudo-code is given below. As a result, *M* ′[*i, j*] is a much more accurate quantitative measure for *i* ∼ *j*, as now it incorporates the equivalent signals from alignments across all units. The algorithm, although runs in *O*(|*S*|^3^) for the worst case, is fairly fast in practice, as *M* is a sparse matrix.

### Algorithm 1

Refine initial matrix *M* to produce matrix *M* ′

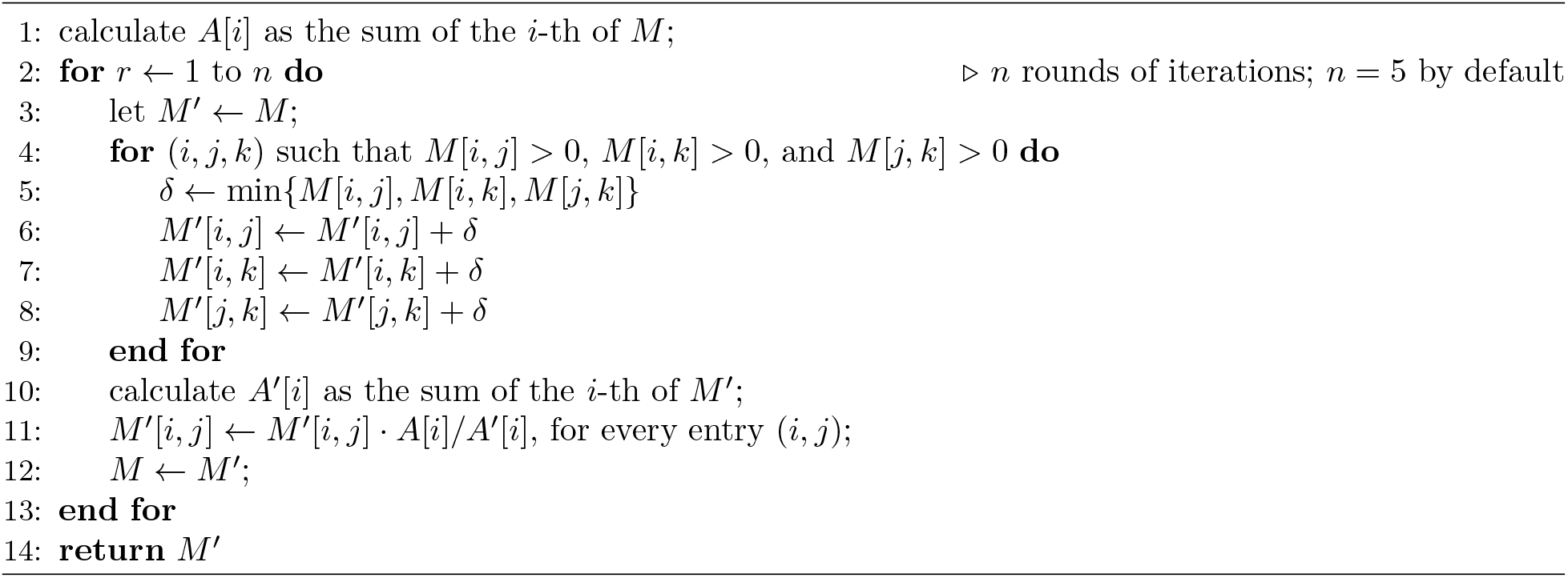

We now extract pairs (*i, j*) where *i* and *j* are likely equivalent. Since equivalent pairs are expected to form paths, we find paths in *M* ′. Specifically, we use a dynamic programming algorithm to iteratively find paths consisting of entries connected vertically, horizontally, or diagonally. The score of a path *P*, denoted as *f* (*P*), is defined as the sum of the values in *M* ′ along the path, i.e., *f* (*P*) = ∑ _(*i,j*)∈*P*_ *M* ′[*i, j*]. We compute all non-overlapping paths with scores at least 1, collected as 𝒫.

We group all |*S*| positions into equivalent classes following paths in 𝒫. We implement this using a disjoint-set data structure. Initially, each position forms its own equivalent class. We collect all entries covered by 𝒫, and sort them according to *M* ′[, ] in descending order. Then for each entry (*i, j*), we combine the two classes including *i* and *j* respectively into a single class. In this process, we impose a restriction that prevents any two positions within a distance of 5 from appearing in the same class. This restriction reduces false combines but may be risky for short unit lengths. To accommodate this we develop a heuristic to handle short repeating units separately. At the end, we obtain a set of equivalent classes, denoted as 𝒞, each of which consists of a set of positions. For each equivalence class, we perform a majority vote to determine the representative letter of that class.

**Step 3: constructing candidate units from 𝒞**. We construct a weighted graph *G* = (𝒞, *E*) where each node represents an equivalent class and edges connect equivalent classes with (nearly) consecutive positions.

Let *A, B* ∈ 𝒞 be two equivalent classes. Let *E*_1_ := {(*i, i* + 1) | *i* ∈ *A, i* + 1 ∈ *B*} be the set of consecutive positions in *A* and *B*. Let *E*_2_ := {(*i, i*+ 2) | *i* ∈ *A, i*+ 2 ∈ *B*, and *i* + 1 forms its own equivalent class} be the set of “gapped” consecutive positions where the gap is likely an insertion. If |*E*_1_| + |*E*_2_| ≥ 1 then edge (*A, B*) is added to *G* and the edge weight *w*(*A, B*) is set as |*E*_1_| + |*E*_2_|. In the constructed graph *G*, we identify the cycle *C* whose smallest edge weight is maximized while ensuring that the equivalent classes on *C* collectively cover more than half of the positions in the repeat region. This can be implemented using binary search combined with BFS or DFS. The cycle will be then transformed into the inferred unit by concatenating the associated letters. We denote by *U*_*_ this inferred unit.

False combinations in constructing 𝒞 may still happen, which often result in incorrect units of shorter length (see Fig. 3). The equivalent classes caused by over-combination are characterized by abnormal size and incoming nodes connecting to different positions. We design a heuristic to split over-combined equivalent classes. We only try to split classes whose size is at least twice the median class size. For each such equivalent class *B*, we examine the in-edge (*A, B*) with maximized weight. Positions in *B* that “follow” positions in *A* will be moved to new class *B*_1_, and others will be moved to another new class *B*_2_. Formally, *B*_1_ := {*i* + 1 ∈ *B* | *i* ∈ *A*} ∪ {*i* + 2 ∈ *B* | *i* ∈ *A* and *i* + 1 forms its own equivalent class}, and *B*_2_ = *B* \ *B*_1_. *B* will be replaced by *B*_1_ and *B*_2_ and edge weights adjacent to *B*_1_ and *B*_2_ will be calculated in the same way above. We iterate over all classes, splitting each oversized class, and repeat this process for up to five rounds. In each round, we reduce the size of the smallest class that can be split by one, while also ensuring it is greater than the median class size. After each round of node splitting, we use the same approach on the new graph to identify a cycle and reconstruct a candidate unit 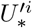 where *i* = 1, 2, · · ·, 5.

**Fig 3:**
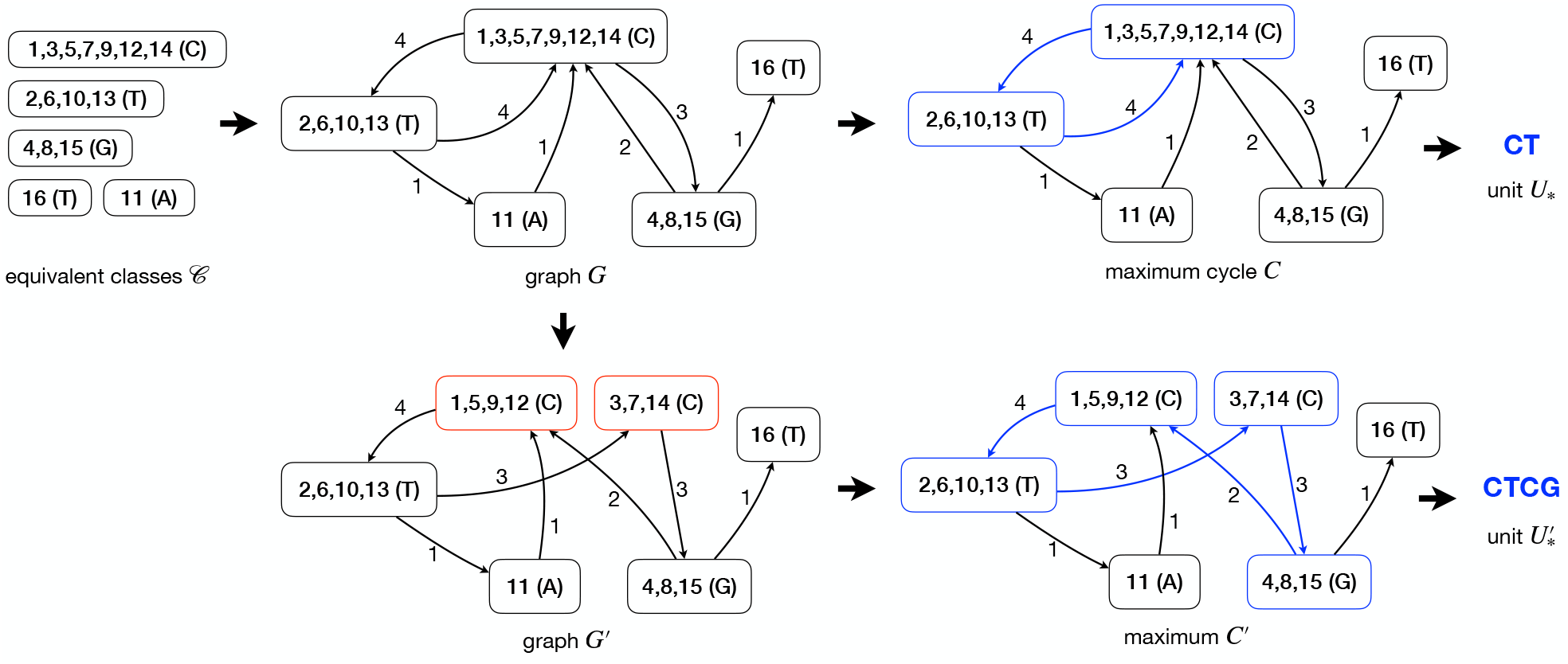
Illustration of constructing the two candidate units *U*_*_ and 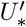, given the equivalent classes 𝒞.

We now describe the heuristic to handle small unit sizes. The core idea is that if the unit is small, some instances of the unit likely remain intact, making the most frequent one a probable candidate for the true unit. Specifically, for each size *k, k* = 2, 3, 4, 5, 6, we identify the most frequent kmer in S by rotating each kmer to its lexicographically smallest form. This most frequent kmer is selected as the candidate unit, denoted as 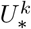. The outcome of Step 3 is 11 candidate units: 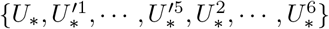.

**Step 4: selecting the optimal unit**. For each candidate unit, we concatenate it multiple times to match the length of |*S*|, and calculate the edit distance between the concatenation and *S*. We choose the unit with the lowest edit distance.

## 3 Results

We implemented the algorithm described in Methods as a new tandem repeat detection tool EquiRep. We compare EquiRep to four other repeat detectors: TRF, mTR, mreps, and TideHunter. For a given input sequence, each of these methods can generate multiple repeat patterns as the output while EquiRep generates a single repeat pattern. If there are multiple predictions, we choose the unit corresponding to a criterion (for example, maximum copy number) best for the method as the final predicted sequence. We evaluate these methods both on simulated and real datasets as follows.

### 3.1 Evaluation with Simulated Random Sequences

The simulated random sequences are generated as follows: 1, generate a random string *U* constituting nucleotides (A,T,G,C) of length 10, 200, and 500, which serves as the ground truth repeat unit; 2, concatenate multiple copies of the unit *U* to generate a longer sequence, with frequency of units being 3, 5, 10, and 20; 3, introduce random errors—insertions, deletions, and substitutions at equal probabilities—at rates of 10% and 20% into the concatenated string to simulate real-world sequencing errors; 4, insert random strings, matching the length of the concatenated string (i.e., the repeat region), at both sides of the concatenated string.

For each of the 40 settings (the combination of unit length, frequency of units, and error rate), we randomly and independently generate 50 sequences. We evaluate the methods’ predictions as follows. Let *T* be a ground-truth repeat unit and let *P* be a prediction. We compute a rotation-aware edit distance between *P* and *T*. Since *P* may be a rotation of the *T*, we calculate the edit distance between *T* and all possible rotations of *P*, and take the minimum value. For each setting, we analyze the 50 instances and report the following metrics: the number of exact matches (i.e., rotation-aware edit distance is 0), the number of instances with a rotation-aware edit distance less than 10% and 20% of the unit length, and the average rotation-aware edit distance across all 50 instances.

Fig. 4 shows the number of correct instances on simulated data at 10% error rate for various lengths and copy numbers. EquiRep consistently predicts a comparable or greater number of correct instances than other methods. The methods with performance closest to EquiRep appear to be mTR and TRF; however, both struggle to maintain accuracy with low copy numbers and very large unit lengths, such as 500 bp. The accuracy of EquiRep is significantly higher than any of the other methods for unit length 500 bp which demonstrates the ability of our tool to predict longer tandem repeats. The number of instances with edits less than 10% of the unit length ranges within 40-50 for EquiRep regardless of the copy number (see Fig. 5) and the trend tends to be consistent over the different unit lengths, unlike other methods. Moreover, EquiRep consistently achieves the lowest average edit distance (see Fig. 6), indicating that even when its predictions are incorrect, they remain close to the true sequence. Results for 20% error rate are available in Supplementary Fig. 1, Supplementary Fig. 2, and Supplementary Fig. 3. For higher error rate, TRF, mreps, and TideHunter see a sharp decline in accuracy as the unit length exceeds 10 bp. Conversely, mTR’s ability to handle long, noisy reads allows it to achieve accuracy close to EquiRep; however, its performance drops when the unit length reaches 500 bp. At a unit length of 500 bp and with high sequencing errors, all methods struggle to accurately predict tandem repeats, particularly when the copy number is low. Overall, EquiRep outperforms every other tool on the three metrics consistently across different simulations.

**Fig 4:**
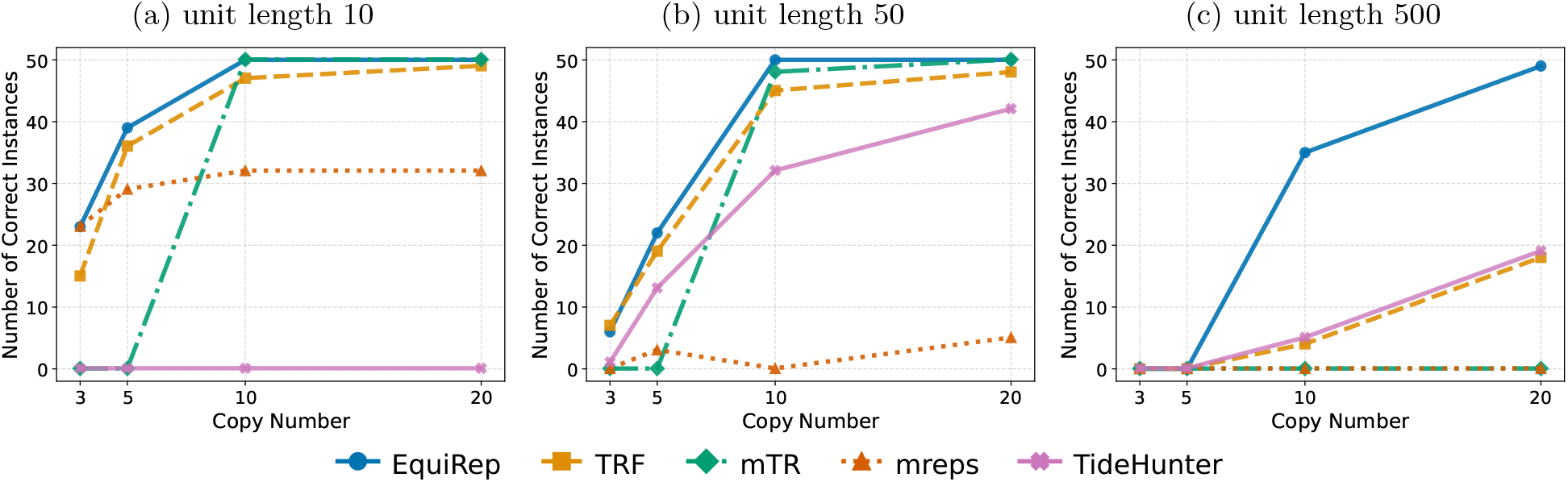
Comparison of number of correct predictions on simulated data at 10% error rate.

**Fig 5:**
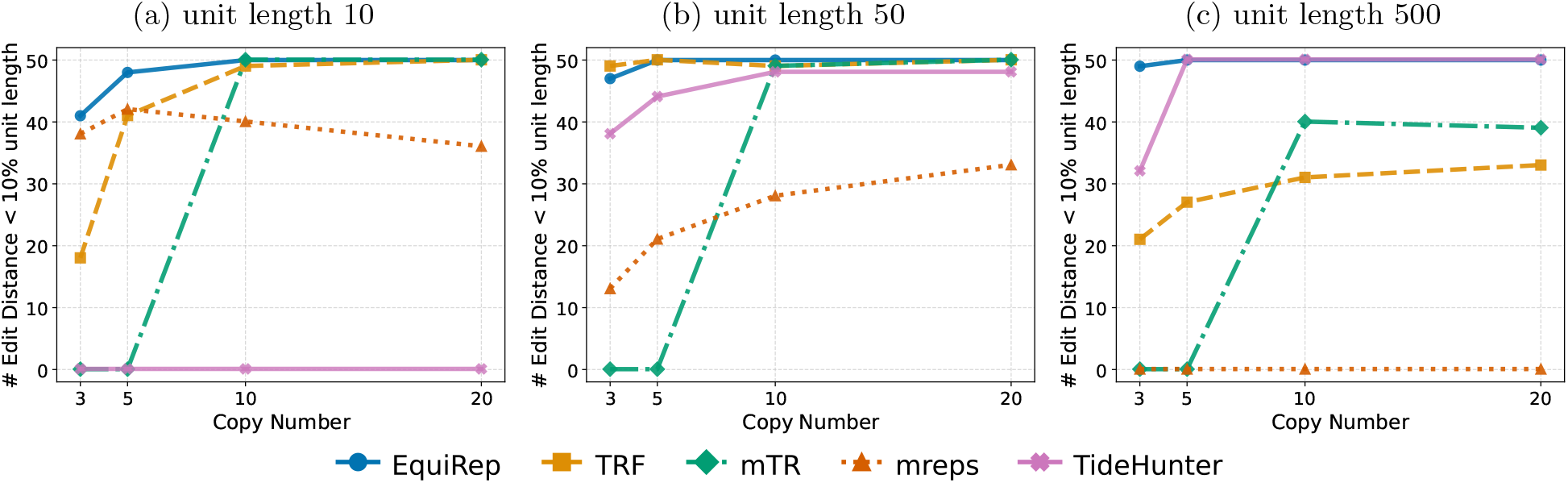
Comparison of number of instances with edits less than 10% of the unit length on simulated data at 10% error rate.

**Fig 6:**
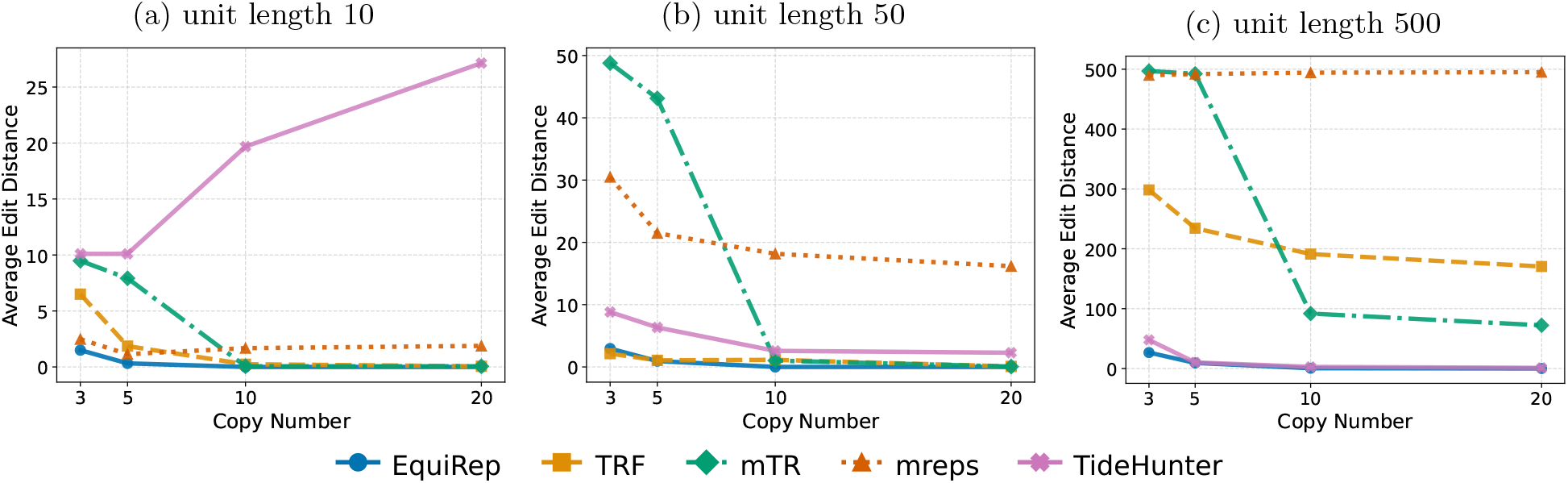
Comparison of average edit distance on simulated data at 10% error rate.

### 3.2 Evaluation using Simulated Sequences with Recurring kmers in a Unit

Genomic sequences are not pure random, often containing recurring substrings. We compare different methods on this scenario with simulations where the repeat unit itself contains recurring structures. In this setting, predicting the correct repeat sequence is challenging as methods may encounters difficulties in distinguishing between such recurring kmers in a single unit and identical kmers across multiple units.

We use this approach to simulate the above sequences. 1, for a given unit length *l* ∈ {50, 200, 500}, we generated a random kmer of length *k* ∈ {5, 10, 20}, respectively. 2, we construct the repeat unit by concatenating the random kmer 2 or 3 times. After these concatenations, any remaining positions within the unit (i.e., *l* − 2*k* for 2 concatenations and *l* − 3*k* for 3 concatenations) will be filled with random nucleotides. 3, we concatenate multiple copies of the repeat unit to generate a longer sequence, with frequency of units being 3, 5, 10, 20. 4, we introduce random errors at rates of 10% and 20%. 5, At the end we insert random strings, matching the length of the concatenated string at both ends.

The same evaluation metrics as in Section 3.1 are also used here. Fig. 7 indicates that the number of correct instances predicted by EquiRep exceeds or is equal to other methods when the simulations have 2 copies of a kmer within the unit at 10% error rate. Fig. 8 shows that almost all the edits predicted by our method are less than 10% of the unit length. Again, EquiRep achieves the lowest average edits as illustrated in Fig. 9. For similar data with 20% error rate, there is a drastic decline in accuracy for all methods except mTR and EquiRep (Supplementary Fig. 4, Supplementary Fig. 5, Supplementary Fig. 6). Nearly all repeat units generated by EquiRep have edits below 10% of the unit length for copy numbers above 10, which highlights the reliability of our predictions specially in challenging erroneous settings. We also tested all methods on another set of data with 3 copies of repeating kmers within the repeat unit. Supplementary Fig. 7, Supplementary Fig. 8, and Supplementary Fig. 9 demonstrates the performance on this dataset for the same three metrics at 10% error rate. Similar results for 20% error rate are presented in Supplementary Fig. 10, Supplementary Fig. 11, and Supplementary Fig. 12. EquiRep is able to make better or similar predictions in all cases indicating that its algorithm is least affected by the presence of embedding kmers within repeat units.

**Fig 7:**
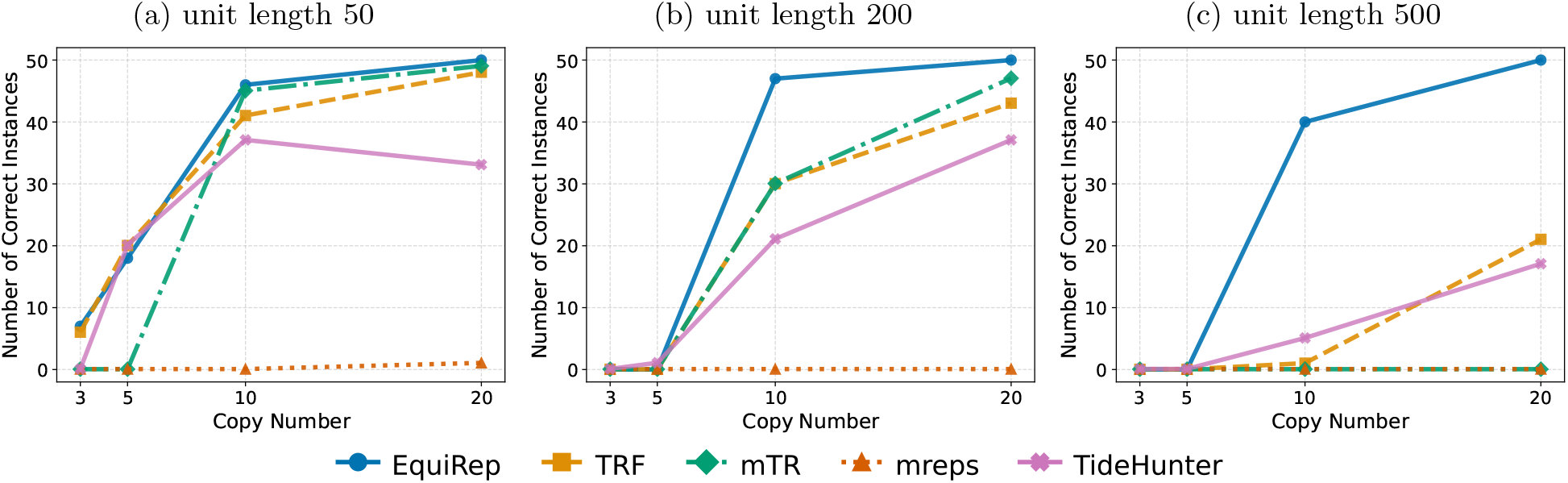
Comparison of number of correct predictions on simulations with 2 recurring kmers at 10% error rate.

**Fig 8:**
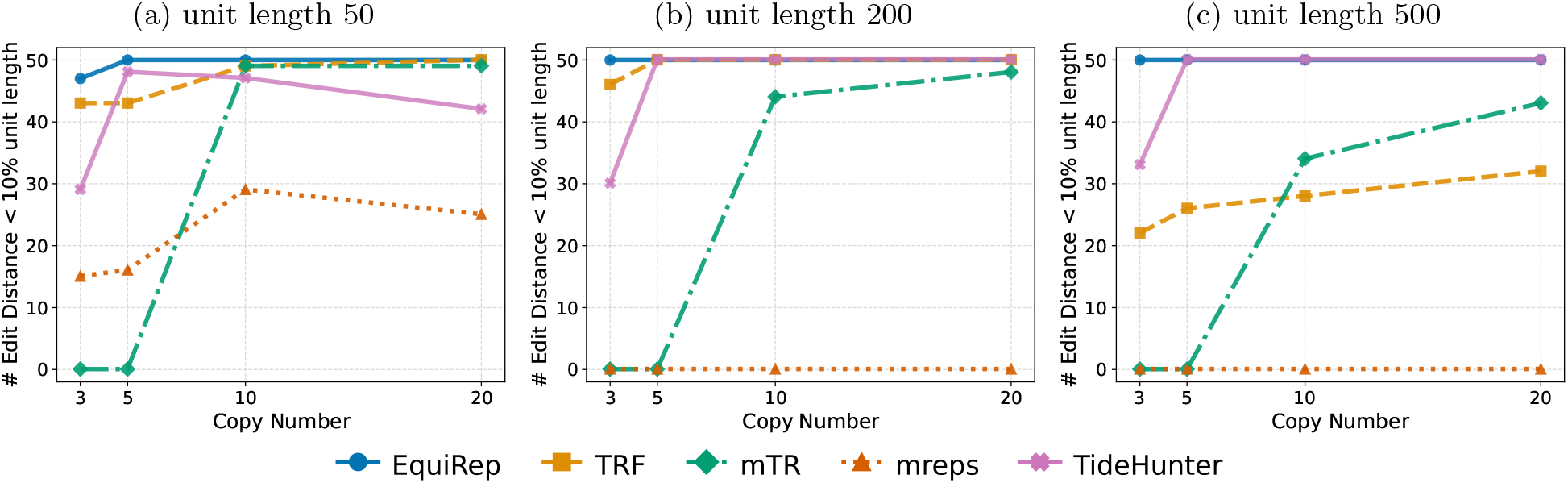
Comparison of number of instances with edits less than 10% of the unit length on simulations with 2 recurring kmers at 10% error rate.

**Fig 9:**
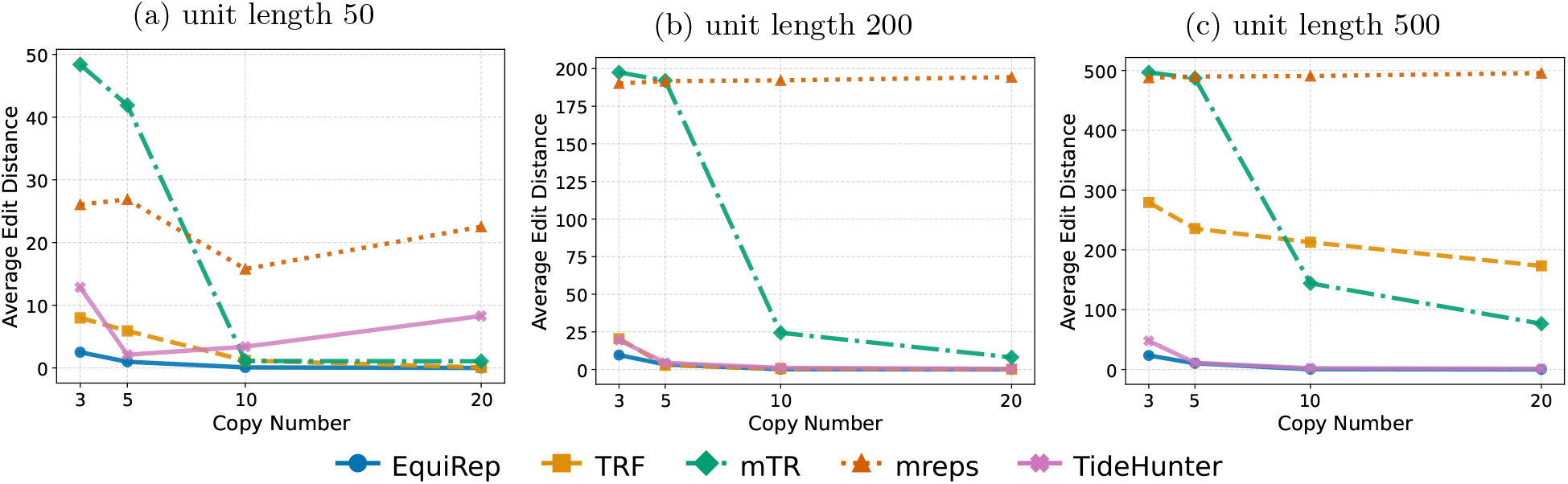
Comparison of average edit distance on simulations with 2 recurring kmers at 10% error rate.

### 3.3 Evaluation using Higher Order Repeat (HOR) Data

We then test all methods on reconstructing unit of a HOR data in human chromosome 5 [16]. This known HOR consists of 13 units, each of which is of size around 171bp. To construct the input sequence for methods to predict, we concatenate the 13 consensus sequences into a string denoted as (x). To create more testing instances, we introduce flanking regions on both sides of the concatenation denoted as (axa), and introduce errors of 1%, 5%, and 10% to (x) and (axa). To evaluate the predicted unit by different methods, we calculate the average rotation-aware edit distance between the predicted unit with each of the 13 known units.

Table 1 shows the results on the HOR data. EquiRep consistently maintains an average edit distance between 21 and 23, outperforming or matching all other tools. The values for EquiRep are similar to mTR when the input sequences have flanking regions at either end (axa) but our method is about 87% better than mTR when just the repeat region is provided (x). Although TideHunter and TRF exhibit accuracy levels similar to ours, they fall short at higher error rates, where EquiRep excels with an 87% improvement.

**Table 1:** Averaged rotation-aware edit distance on HOR data.

| Error Rate (%) | Pattern | EquiRep | mTR | TRF | mreps | TideHunter |
| --- | --- | --- | --- | --- | --- | --- |
| 0 | **x** | 21.54 | 170.31 | 21.77 | 162.31 | 22.31 |
| 0 | **axa** | 21.46 | 21.54 | 21.77 | 166.31 | 22.31 |
| 1 | **x** | 21.38 | 170.31 | 24.08 | 162.31 | 21.92 |
| 1 | **axa** | 21.38 | 21.54 | 24.08 | 162.31 | 21.92 |
| 5 | **x** | 21.92 | 168.31 | 38.77 | 157.31 | 25.46 |
| 5 | **axa** | 21.69 | 21.62 | 38.77 | 158.31 | 25.46 |
| 10 | **x** | 23.31 | 170.31 | 170.31 | 160.31 | 180.39 |
| 10 | **axa** | 21.38 | 25.62 | 170.31 | 165.31 | 180.39 |

### 3.4 Evaluation with Rolling Circle Amplification (RCA) Data

The second set of real data is a RCA based Nanopore sequencing protocol from isocirc [23] that has been used to detect a catalogue of full-length circular RNAs from 12 human tissues. We consider a subset of 101 sequences of the Nanopore long reads from the prostate tissue (GEO accession number: GSE141693) for analysis. It is difficult to evaluate the repeats from the RCA based long reads data due to lack of reliable ground truth, so we evaluate this data in two different ways. Firstly, we use a dot plot analysis. Dot plots have served as a common approach for visualizing and identifying the structural patterns of sequences such as repeats. We first align the input sequence to itself with LASTZ [8] using specific parameters designed for generating dot plots. The alignment program generates a dot file which can be converted to an image file for visualization using a simple R script. The dot file can be used to estimate the repeat unit length. We treat this estimate as a benchmark for comparing the predictions of EquiRep and other tools. We report the number of predictions that fall within 5%, 20%, 50%, and 80% error range of the true length. For the second approach, we first concatenate copies of the unit predicted to get a string A which is longer than the input sequence. Then we get the “semi-edit distance” which is the smallest edit distance between any substring of A and the input sequence. We record it and report the number of predictions that has a ratio (semi-edit-distance)/(input sequence-length) less than or equal to 0.1, 0.2, 0.3, 0.5, 0.8.

Table 2 compares different methods in terms of the predicted repeat unit length, and Table 3 compares the normalized semi-edit-distance. In both metrics, EquiRep demonstrates high accuracy, consistently out-performing mTR, TRF, and mreps. The results are also comparable to TideHunter, which is specifically optimized for RCA-based analysis. Given that the exact repeat sequences for this dataset are not available, similar metric values in the table can be interpreted as comparable accuracy. It should be noted that while TideHunter excels on RCA data, its accuracy diminishes on shorter unit repeats as indicated by the simulation results. This highlights that EquiRep is adaptable to a broad range of complex sequences and versatile for various applications.

**Table 2:** Performance on RCA data: number of predicted repeat lengths within error ranges of the true length and number of no repeats found (out of 101 instances).

| Error Range | EquiRep | mTR | TRF | mreps | TideHunter |
| --- | --- | --- | --- | --- | --- |
| 0.95 to 1.05 (5%) | 98 | 5 | 68 | 1 | 101 |
| 0.8 to 1.2 (20%) | 100 | 5 | 68 | 1 | 101 |
| 0.5 to 1.5 (50%) | 100 | 5 | 68 | 1 | 101 |
| 0.2 to 1.8 (80%) | 101 | 9 | 69 | 1 | 101 |
| #norepeat | 0 | 18 | 30 | 7 | 0 |

**Table 3:** Performance on RCA data: number of predicted repeat units with ratio of edit distance to input length less than various percentages (out of 101 instances). SED = semi-edit-distance.

| SED/Length | EquiRep | mTR | TRF | mreps | TideHunter |
| --- | --- | --- | --- | --- | --- |
| $\leq 0.05$ (5%) | 0 | 0 | 0 | 0 | 0 |
| $\leq 0.1$ (10%) | 67 | 5 | 52 | 0 | 73 |
| $\leq 0.2$ (20%) | 99 | 5 | 68 | 1 | 101 |
| $\leq 0.3$ (30%) | 101 | 5 | 68 | 1 | 101 |
| $\leq 0.5$ (50%) | 101 | 28 | 69 | 40 | 101 |
| $\leq 0.8$ (80%) | 101 | 83 | 71 | 94 | 101 |

### 3.5 Comparison of Running Time

Supplementary Table 1 shows the runtime in seconds for all methods including EquiRep on simulated data. TideHunter and TRF has the fastest average execution times. Although EquiRep has a longer execution time compared to the other methods, it is acceptable and at the similar level with mTR. Moreover, there is potential for huge efficiency improvements (see Discussion). EquiRep’s superior accuracy compensates for its longer runtime, reinforcing its overall effectiveness in tandem repeat detection.

## 4 Conclusion and Discussion

Tandem repeats play critical roles in genetic diversity, gene regulation, and have strong associations with various neurological and developmental disorders. Accurate detection of tandem repeats is essential for its analysis, but yet an unsolved problem due to complicated scenarios of repeating unit and highly erroneous sequences. In this paper, we present EquiRep, a robust and accurate tool for repeat detection. By leveraging a unique approach of grouping nucleotide positions into equivalent classes, EquiRep effectively builds a weighted graph to reconstruct repeat units with high accuracy. Our method addresses key challenges in detecting both short and long tandem repeats from highly erroneous sequences, areas where existing tools often fall short. Through extensive testing on both simulated and real HOR and RCA datasets, EquiRep outperforms or matches current state-of-the-art methods, demonstrating its robustness to sequencing errors and complex repeat patterns.

We are optimistic that the computational efficiency of EquiRep can be largely improved. Currently, the self-local alignment step presents a bottleneck in runtime. By improving this step, possibly through adapting more efficient alignment algorithms or parallel processing, we can substantially reduce its runtime. The second time-consuming step in EquiRep is matrix refinement. Matrix operations are inherently parallelizable, and the sparse property of the matrix can be leveraged to achieve an acceleration. We plan to explore these approaches to improve the runtime of EquiRep.

We also aim for improving EquiRep’s accuracy, particularly for RCA data. The framework of EquiRep allows it to be improved in several ways. One approach is to enhance matrix refinement, which is crucial for producing accurate equivalent classes. The current method considers three mutually supportive pairs, but it can be extended to account for insertions and deletions. More precise modeling of insertions and deletions using equivalent classes, rather than single positions, is expected to improve node splitting, a key step in rescuing over-combines. Initial predictions of unit length might also help with guiding the search for repeat units within a specified range. Finally, improved heuristics for identifying cycles that combine both weights and optimal positional coverage would enable the weighted graph to represent complex repeat patterns more accurately. We plan to explore these strategies to enhance EquiRep’s accuracy.

## Availability

The source code of EquiRep is freely available at https://github.com/Shao-Group/EquiRep. The scripts, evaluation pipelines, and instructions that can be followed to reproduce the experimental results of this work is available at https://github.com/Shao-Group/EquiRep-test. The Nanopore long reads data (fastq) from the prostate sample is available at GEO (accession number: GSE141693).

## Acknowledgment

This work is supported by the US National Science Foundation (2145171 to M.S.) and by the US National Institutes of Health (R01HG011065 to M.S.).

## References

1. Benson, G.: Tandem repeats finder: a program to analyze dna sequences. Nucleic acids research 27(2), 573–580 (1999)

2. Campuzano, V., Montermini, L., Molto, M.D., Pianese, L., Cossée, M., Cavalcanti, F., Monros, E., Rodius, F., Duclos, F., Monticelli, A., et al.: Friedreich’s ataxia: autosomal recessive disease caused by an intronic gaa triplet repeat expansion. Science 271(5254), 1423–1427 (1996)

3. De Roeck, A., Duchateau, L., Van Dongen, J., Cacace, R., Bjerke, M., Van den Bossche, T., Cras, P., Vandenberghe, R., De Deyn, P.P., Engelborghs, S., et al.: An intronic vntr affects splicing of abca7 and increases risk of alzheimer’s disease. Acta neuropathologica 135, 827–837 (2018)

4. Dolzhenko, E., Deshpande, V., Schlesinger, F., Krusche, P., Petrovski, R., Chen, S., Emig-Agius, D., Gross, A., Narzisi, G., Bowman, B., et al.: Expansionhunter: a sequence-graph-based tool to analyze variation in short tandem repeat regions. Bioinformatics 35(22), 4754–4756 (2019)

5. Fang, L., Liu, Q., Monteys, A.M., Gonzalez-Alegre, P., Davidson, B.L., Wang, K.: Deeprepeat: direct quantification of short tandem repeats on signal data from nanopore sequencing. Genome biology 23(1), 108 (2022)

6. Gao, Y., Liu, B., Wang, Y., Xing, Y.: Tidehunter: efficient and sensitive tandem repeat detection from noisy long-reads using seed-and-chain. Bioinformatics 35(14), i200–i207 (2019)

7. Hannan, A.J.: Tandem repeats mediating genetic plasticity in health and disease. Nature Reviews Genetics 19(5), 286–298 (2018)

8. Harris, R.S.: Improved pairwise alignment of genomic DNA. The Pennsylvania State University (2007)

9. Kolpakov, R., Bana, G., Kucherov, G.: mreps: efficient and flexible detection of tandem repeats in dna. Nucleic acids research 31(13), 3672–3678 (2003)

10. Kristensen, L.S., Jakobsen, T., Hager, H., Kjems, J.: The emerging roles of circRNAs in cancer and oncology. Nature Reviews Clinical Oncology 19(3), 188–206 (2022)

11. Liu, Z., Tao, C., Li, S., Du, M., Bai, Y., Hu, X., Li, Y., Chen, J., Yang, E.: circfl-seq reveals full-length circular rnas with rolling circular reverse transcription and nanopore sequencing. elife 10, e69457 (2021)

12. Logsdon, G.A., Rozanski, A.N., Ryabov, F., Potapova, T., Shepelev, V.A., Catacchio, C.R., Porubsky, D., Mao, Y., Yoo, D., Rautiainen, M., et al.: The variation and evolution of complete human centromeres. Nature 629(8010), 136–145 (2024)

13. Melters, D.P., Bradnam, K.R., Young, H.A., Telis, N., May, M.R., Ruby, J.G., Sebra, R., Peluso, P., Eid, J., Rank, D., et al.: Comparative analysis of tandem repeats from hundreds of species reveals unique insights into centromere evolution. Genome biology 14, 1–20 (2013)

14. Mitsuhashi, S., Frith, M.C., Mizuguchi, T., Miyatake, S., Toyota, T., Adachi, H., Oma, Y., Kino, Y., Mitsuhashi, H., Matsumoto, N.: Tandem-genotypes: robust detection of tandem repeat expansions from long dna reads. Genome biology 20, 1–17 (2019)

15. Morishita, S., Ichikawa, K., Myers, E.W.: Finding long tandem repeats in long noisy reads. Bioinformatics 37(5), 612–621 (2021)

16. Paar, V., Basar, I., Rosandic, M., Gluncic, M.: Consensus higher order repeats and frequency of string distributions in human genome. Current genomics 8(2), 93–111 (2007)

17. Rybak-Wolf, A., Stottmeister, C., Glažar, P., Jens, M., Pino, N., Giusti, S., Hanan, M., Behm, M., Bartok, O., Ashwal-Fluss, R., Herzog, M., Schreyer, L., Papavasileiou, P., Ivanov, A., Öhman, M., Refojo, D., Kadener, S., Rajewsky, N.: Circular RNAs in the Mammalian Brain Are Highly Abundant, Conserved, and Dynamically Expressed. Molecular Cell 58(5), 870–885 (2015)

18. Siwach, P., Ganesh, S.: Tandem repeats in human disorders: mechanisms and evolution. Front. Biosci 13, 4467–4484 (2008)

19. Song, J.H., Lowe, C.B., Kingsley, D.M.: Characterization of a human-specific tandem repeat associated with bipolar disorder and schizophrenia. The American Journal of Human Genetics 103(3), 421–430 (2018)

20. Usdin, K.: The biological effects of simple tandem repeats: lessons from the repeat expansion diseases. Genome research 18(7), 1011–1019 (2008)

21. Wang, F., Nazarali, A.J., Ji, S.: Circular RNAs as potential biomarkers for cancer diagnosis and therapy. American Journal of Cancer Research 6(6), 1167–1176 (2016)

22. Wirawan, A., Kwoh, C.K., Hsu, L.Y., Koh, T.H.: Inverter: integrated variable number tandem repeat finder. In: Computational Systems-Biology and Bioinformatics: First International Conference, CSBio 2010, Bangkok, Thailand, November 3-5, 2010. Proceedings. pp. 151–164. Springer (2010)

23. Xin, R., Gao, Y., Gao, Y., Wang, R., Kadash-Edmondson, K.E., Liu, B., Wang, Y., Lin, L., Xing, Y.: isoCirc catalogs full-length circular RNA isoforms in human transcriptomes. Nature communications 12(1), 266 (2021)

24. Zhang, J., Hou, L., Zuo, Z., Ji, P., Zhang, X., Xue, Y., Zhao, F.: Comprehensive profiling of circular rnas with nanopore sequencing and ciri-long. Nature biotechnology 39(7), 836–845 (2021)

