## Supplementary Material for "Accurate Detection of Tandem Repeats from Error-Prone Sequences with EquiRep"

### Supplementary Materials for "Accurate Detection of Tandem Repeats from Error-Prone Sequences with EquiRep"

#### List of Supplementary Tables

1 Comparison of running time in seconds fro different lengths and copy numbers on simulated data. 6

†These authors contributed equally to this work.

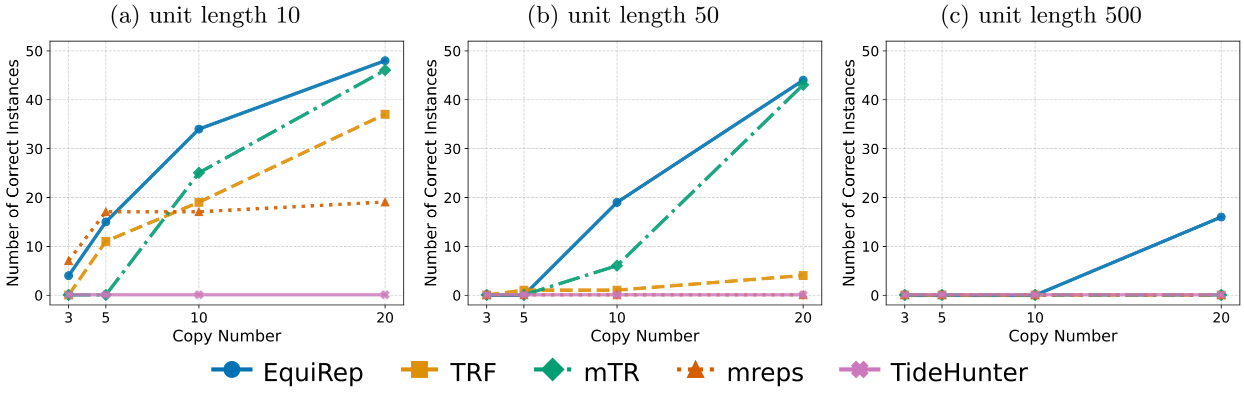

**Supplementary Figure 1**: Comparison of number of correct predictions on simulated data at 20% error rate.

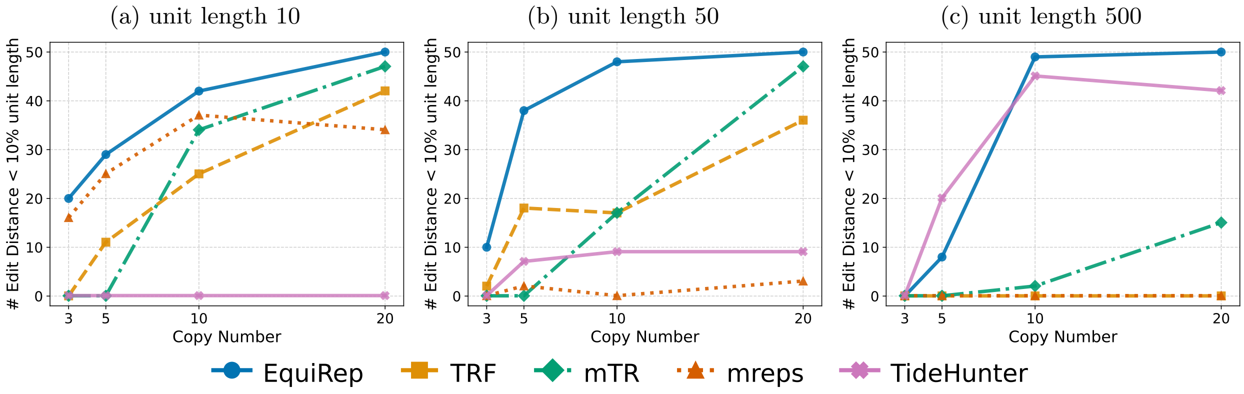

**Supplementary Figure 2**: Comparison of number of instances with edits less than 10% of the unit length on simulated data at 20% error rate.

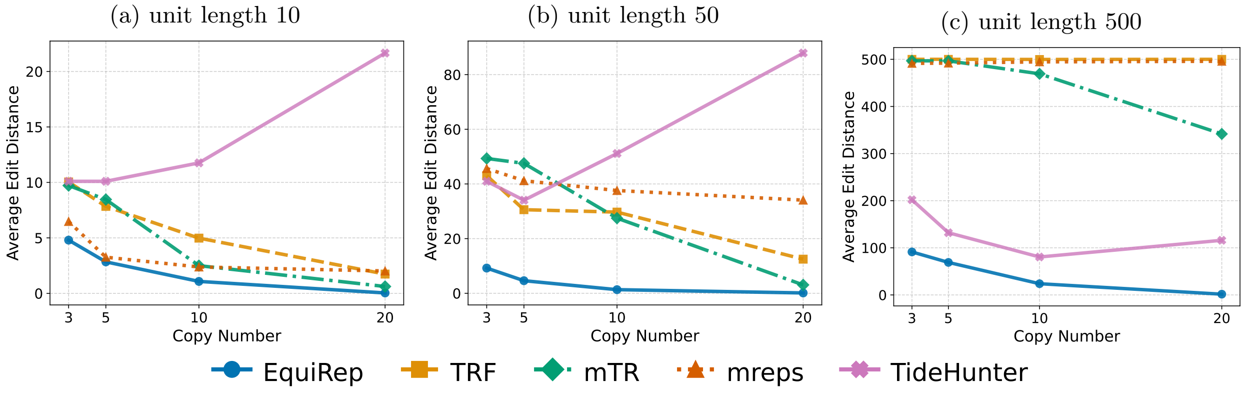

**Supplementary Figure 3**: Comparison of average edit distance on simulated data at 20% error rate.

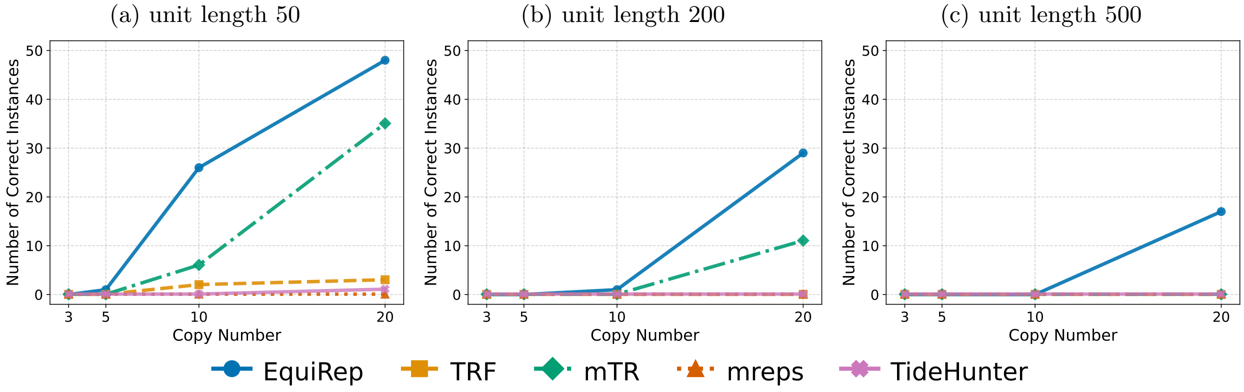

**Supplementary Figure 4**: Comparison of number of correct predictions on simulations with 2 recurring kmers at 20% error rate.

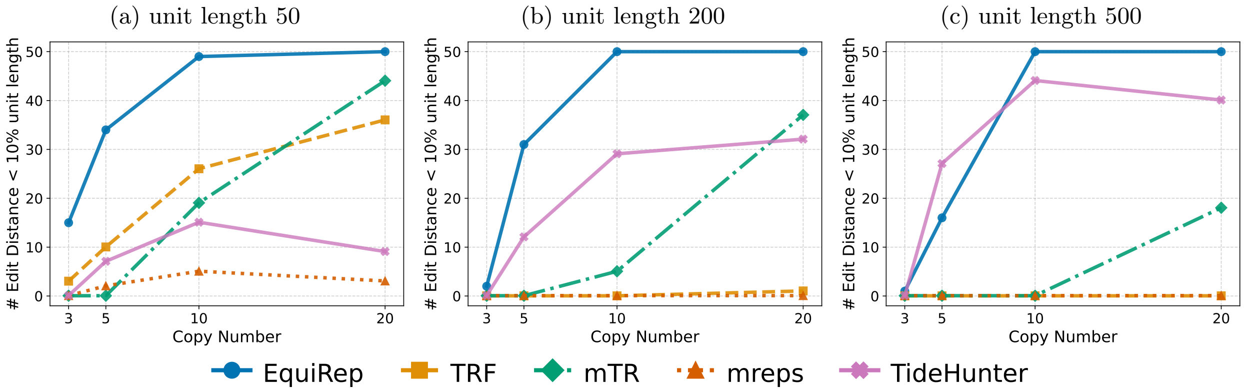

**Supplementary Figure 5**: Comparison of number of instances with edits less than 10% of the unit length on simulations with 2 recurring kmers at 20% error rate.

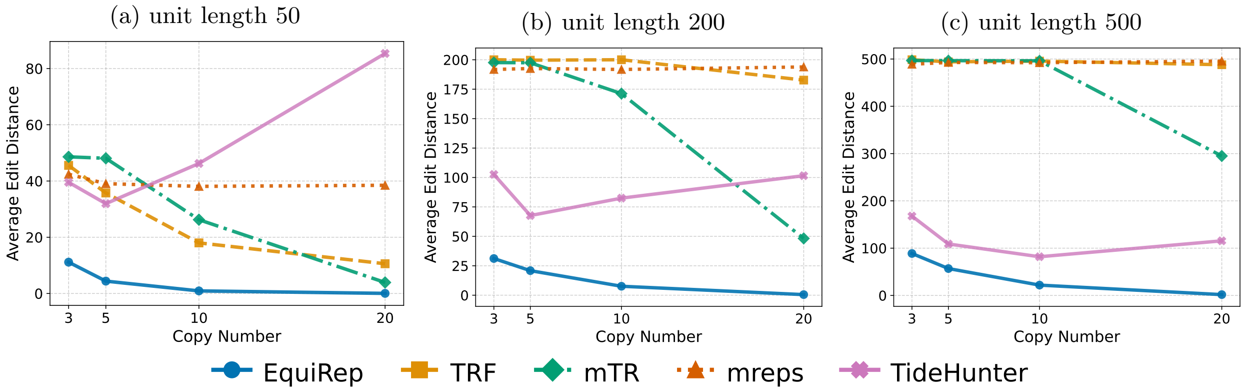

**Supplementary Figure 6**: Comparison of average edit distance on simulations with 2 recurring kmers at 20% error rate.

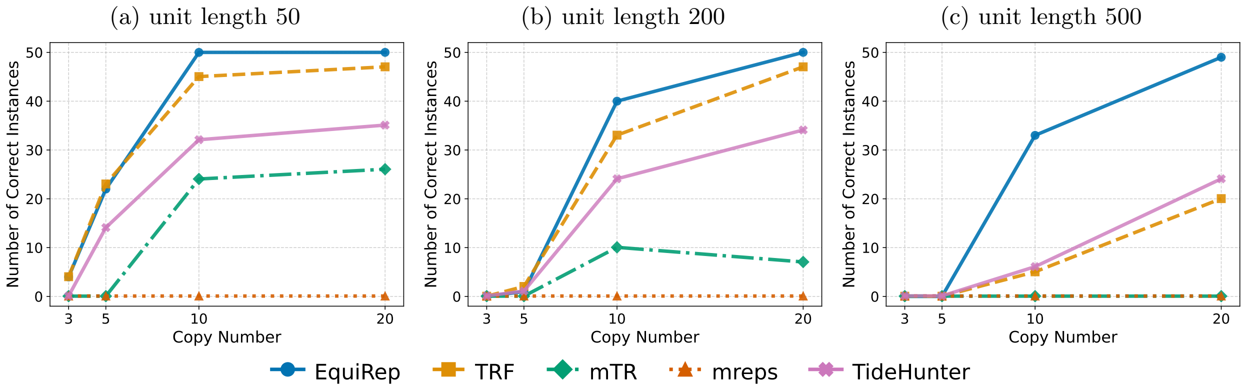

**Supplementary Figure 7**: Comparison of number of correct predictions on simulations with 3 recurring kmers at 10% error rate.

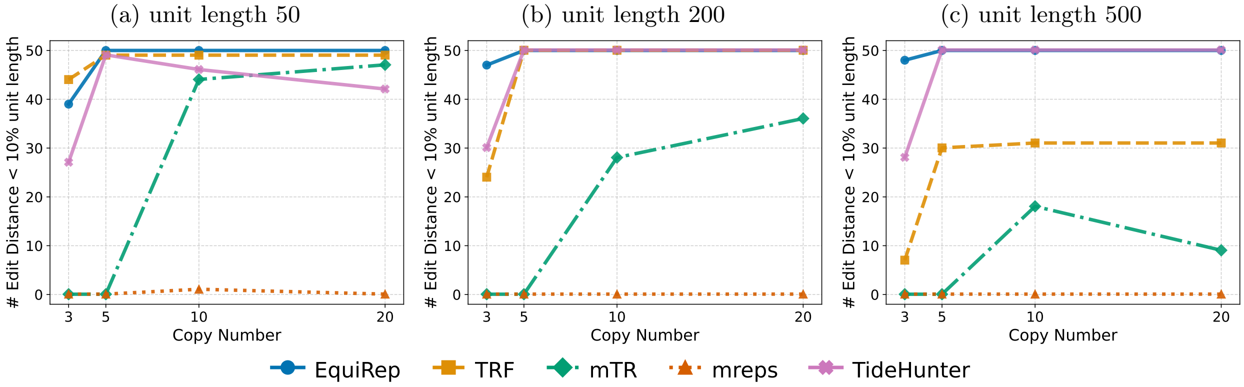

**Supplementary Figure 8**: Comparison of number of instances with edits less than 10% of the unit length on simulations with 3 recurring kmers at 10% error rate.

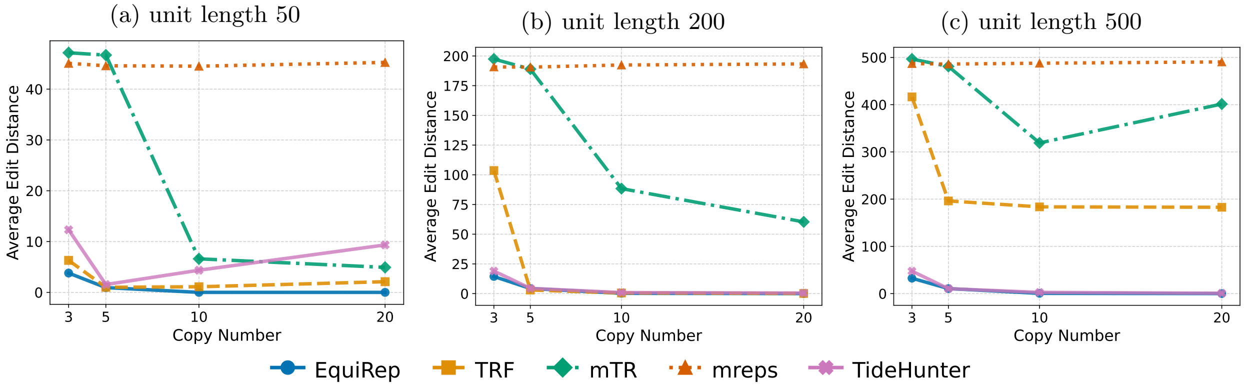

**Supplementary Figure 9**: Comparison of average edit distance on simulations with 3 recurring kmers at 10% error rate.

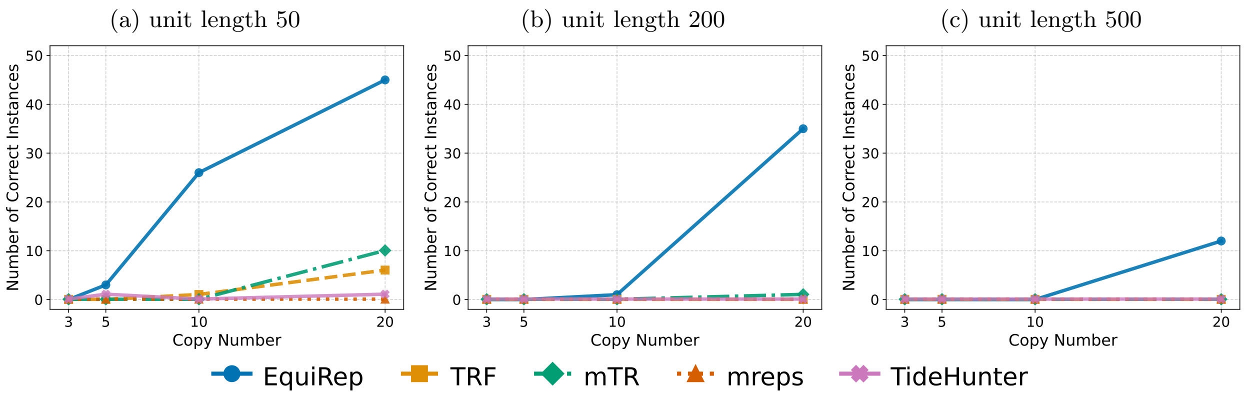

**Supplementary Figure 10**: Comparison of number of correct predictions on simulations with 3 recurring kmers at 20% error rate.

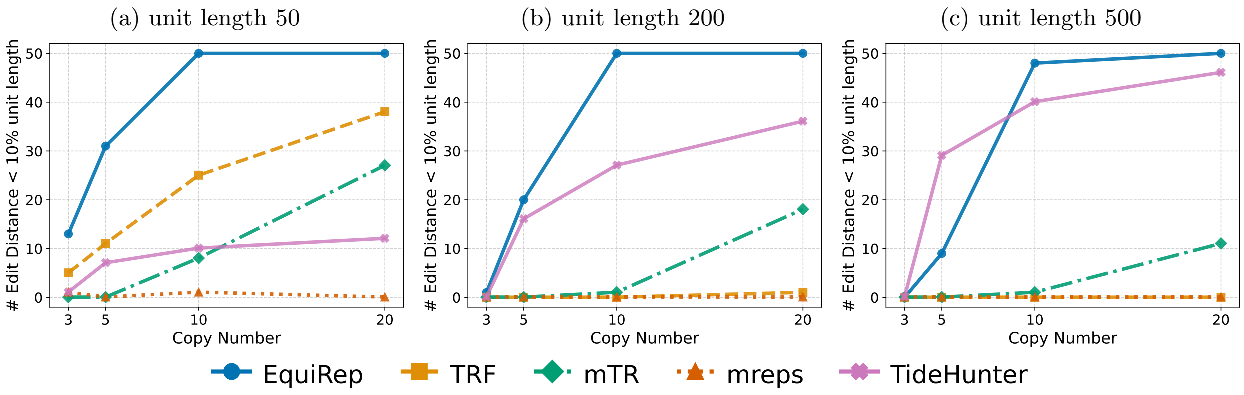

**Supplementary Figure 11**: Comparison of number of instances with edits less than 10% of the unit length on simulations with 3 recurring kmers at 20% error rate.

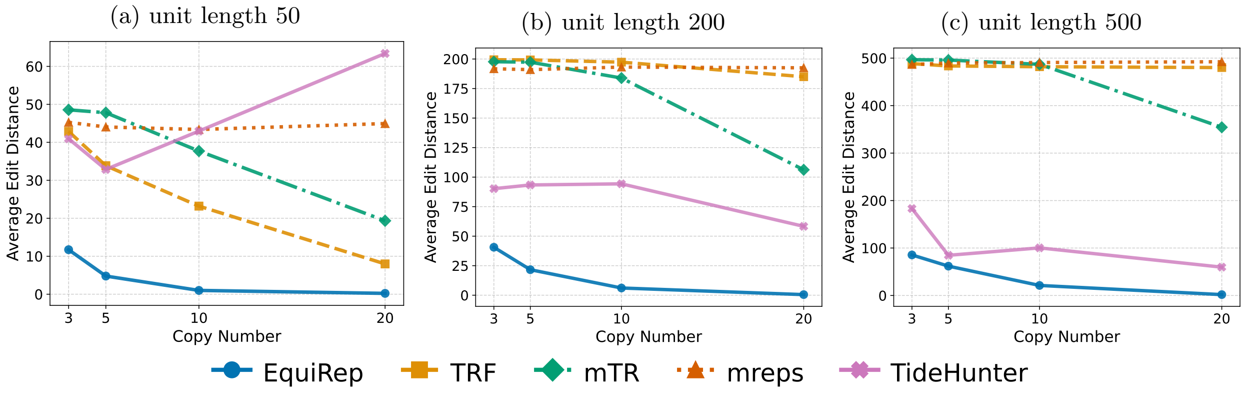

**Supplementary Figure 12**: Comparison of average edit distance on simulations with 3 recurring kmers at 20% error rate.

**Supplementary Table 1**: Comparison of running time in seconds fro different lengths and copy numbers on simulated data.

| **Length_CopyNumber** | **EquiRep** | **TRF** | **mTR** | **mreps** | **TideHunter** |
|---|---|---|---|---|---|
| 5_3 | 3.67 | 0.00 | 1.15 | 0.01 | 0.01 |
| 5_5 | 13.73 | 0.01 | 1.14 | 0.01 | 0.00 |
| 5_10 | 16.18 | 0.03 | 1.04 | 0.02 | 0.02 |
| 5_20 | 14.22 | 0.10 | 1.2 | 0.04 | 0.06 |
| 10_3 | 15.8 | 0.00 | 1.16 | 0.01 | 0.00 |
| 10_5 | 14.18 | 0.03 | 1.04 | 0.02 | 0.02 |
| 10_10 | 14.16 | 0.08 | 1.27 | 0.03 | 0.06 |
| 10_20 | 14.22 | 0.23 | 1.47 | 0.06 | 0.08 |
| 50_3 | 16.24 | 0.04 | 1.4 | 0.10 | 0.04 |
| 50_5 | 14.48 | 0.11 | 1.47 | 0.16 | 0.07 |
| 50_10 | 16.6 | 0.41 | 3.27 | 0.34 | 0.14 |
| 50_20 | 22.97 | 1.32 | 4.59 | 0.69 | 0.19 |
| 200_3 | 16.25 | 0.15 | 2.15 | 0.70 | 0.11 |
| 200_5 | 22.71 | 0.58 | 3.06 | 1.26 | 0.19 |
| 200_10 | 29.98 | 1.51 | 35.67 | 2.67 | 0.33 |
| 200_20 | 74.40 | 3.85 | 45.91 | 5.63 | 1.13 |
| 500_3 | 23.83 | 0.37 | 4.00 | 2.54 | 0.28 |
| 500_5 | 35.07 | 0.85 | 6.11 | 4.47 | 0.57 |
| 500_10 | 104.89 | 2.39 | 90.97 | 9.43 | 2.62 |
| 500_20 | 354.65 | 6.64 | 84.75 | 19.8 | 9.85 |
| **Average** | 41.9115 | 0.935 | 14.641 | 2.3995 | 0.7885 |
